# Selection of sheep skin bacteria to reduce blood-feeding by biting midges under laboratory conditions

**DOI:** 10.1101/2024.05.30.596646

**Authors:** Paula S. Brok, Stéphanie M. Jost, Niels O. Verhulst

## Abstract

Biting midges of the genus Culicoides (Diptera: Ceratopogonidae) are of huge veterinary importance, mainly as vectors of pathogens, such as *Bluetongue virus*. Currently, there are no effective methods to protect animals against biting midges since insecticides have limited or short-lived efficacy. Biting midges are attracted to hosts by carbon dioxide and by their body odours, which are mainly produced by skin bacteria. In humans, it has been shown that differences in attractiveness between individuals to mosquitoes is mediated by these skin bacterial volatiles. This opens the possibility to protect individuals from biting insects by supplementing their skin microbiome with probiotics. In this study, we investigated this approach by culturing sheep skin bacteria on different media and assessed their effects against field-caught Culicoides (overwhelmingly Obsoletus group species) as well as laboratory-reared *Culicoides nubeculosus* (Meigen). *Aerococcus urinaeeequi*, *Bacillus safensis*, *Bacillus subtilis*, *Jeotgalicoccus psychrophilus*, *Micrococcus* sp. and *Staphylococcus equorum* were selected to be tested in a dual-choice Y-tube olfactometer, assessing their behavioural effects towards biting midges. We revealed an avoidance effect towards laboratory-reared *C. nubeculosus* when testing *B. safensis* (P ≤ 0.001) and *B. subtilis* (P ≤ 0.001). *Bacillus safensis* (P = 0.006) and *Micrococcus* sp. (P ≤ 0.001) yielded significant repellent potential towards field-caught Culicoides. These two candidates were subsequently tested in a membrane blood feeding assay. When the bacterial species *B. safensis* was applied to the membrane, a feeding reduction of 83 % was observed with field-caught Culicoides.

## Introduction

Biting midges are of significant veterinary importance, mainly as vectors of pathogens like *Bluetongue virus* (BTV), *Epizootic haemorrhagic disease virus* and *African horse sickness virus*. Further, they cause allergic dermatitis in equids and can be of considerable nuisance also for humans.

Recently, a BTV outbreak in the Netherlands resulted in substantial economic impacts from direct costs due to production losses of the diseased ruminants as well as indirect expenses caused by surveillance programs and animal export restrictions (Stokstad, 2023). The identification of the novel, fast-spreading virus strain BTV-3 in Dutch livestock is particularly alarming, given that currently there is no vaccine available for this strain in Europe.

At present, there are no effective control methods against Culicoides (Harrup et al., 2016; Shults et al., 2021). Establishing physical barriers using nets or fly sheets is impractical for farm animals and proves ineffective due to the small size of biting midges. Insecticides on animals show limited and/or short-lived effectiveness against biting midges, necessitating daily application (Harrup et al., 2016; Venail et al., 2011). Consequently, there is a need for viable and alternative approaches to protect animals against these insects.

Like mosquitoes, biting midges are attracted to their host by carbon dioxide released in their breath and by the body odours the hosts emit (Lucas-Barbosa et al., 2023; Zimmer et al., 2015). These body odours are mainly produced by skin bacteria (Jha, 2017) and mediate the differences in attractiveness between humans to mosquitoes, explaining why some people are bitten more often than others (Showering et al., 2022; Verhulst et al., 2011). Little is known about the odour profiles and the associated skin microbiome of animals, though volatiles from cow and chicken skin bacteria were shown to mediate mosquitoes to their animal host as they significantly increased trap catches (Busula et al., 2017). Recently, the skin bacterial profiles of sheep from three different breeds, farms and body sites were investigated by metabarcoding, revealing that environment rather than breed or body site shapes the skin bacterial community (Jost et al., 2023). We assume that sheep also differ in their attractiveness to biting midges based on the volatiles released by their different skin microbiomes.

Targeting the skin microbiome to inhibit feeding by biting midges could be a novel, long-lasting control tool (Lucas-Barbosa et al., 2022). Several studies on the human skin have shown the beneficial application of skin probiotics to remedy skin dryness or skin diseases such as acne vulgaris and psoriasis (Knackstedt et al., 2020; Yu et al., 2020). Further, it was shown, that the application of lactobacilli can decrease malodour-producing bacteria of human armpits (Onwuliri et al., 2021). It remains to be investigated whether the application of skin probiotics and the associated changes in the odour profile could also reduce feeding by hematophagous insects.

Our goal was to select sheep skin bacterial species that could be used in future studies as a skin probiotic to impair feeding by biting midges. To this aim, we cultured sheep skin bacteria on different media and examined bacterial strains in a Y-tube olfactometer regarding attractiveness or repellency to field-caught and laboratory-reared Culicoides. The bacterial candidates with the highest repellent potential were subsequently investigated in a blood feeding assay.

## Materials and methods

### Skin bacterial samples

Skin bacterial samples were taken as described in Jost et al. (2023) from the back of two adult female sheep (*Ovis aries*; Dorper sheep and Swiss White Alpine sheep) with outdoor access (approved by the cantonal veterinary office; number: 33293). In short, from each sheep, four skin bacteria samples were taken with a dry swab (flocked swab, without diluent; Copan Diagnostics, USA) and another four with a swab in Amies medium (Snappable PS + viscose; Deltalab, Spain) resulting in a total of 16 swabs. The sterile swabs were rubbed ten times over approximately 10 cm of the sheep’s backline, where the wool was parted with fingers (gloves worn). All samples were transported in a polystyrene foam box on dry ice to a freezer and then stored at -20 °C for 3-6 months until further use.

### Skin bacteria cultures

Skin bacterial swabs of each kind and each sheep (eight in total) were used to start cultures in two ml tryptic soy broth (TSB; Sigma-Aldrich, Germany) and in two ml liquid sweat medium. Media were prepared as described before (Lucas-Barbosa et al., 2023). The swabs were manually turned ten times in the respective media which subsequently were incubated at 37 °C for 24 h. Of each resulting culture, 100 μl was spread on plates of Columbia 5 % sheep blood agar (Becton Dickinson, USA), CNA 5 % sheep blood agar (Becton Dickinson), tryptic soy agar (TSA) and sweat media agar.

Additionally, the remaining eight swabs were directly, without cultivation in liquid media, spread onto tryptic soy agar (TSA) and sweat medium agar plates as described (Lucas-Barbosa et al., 2023). All plates were incubated at 37 °C for 24 h. Glycerol stocks were made by mixing 500 μl of the TSB media cultures with 500 μl 50 % glycerol and stored at -80 °C.

Bacterial growth curves were determined by incubating 200 μl (from 100 μl of glycerol stock in 2 ml TSB) of the bacterial sample at 37 °C (except *Jeotgalicoccus psychrophilus* at 24 °C as fortuitously determined) in a 96 well plate in a plate reader (Tecan Infinite 200 PRO, Tecan, Switzerland). Measurements were taken at 10-minute intervals with 8.5 minutes of continuous shaking, 10 seconds of vigorous shaking, and absorbance recorded at 600 nm. Final growth curves were based on at least two technical replicates (Figure S1).

Concentrations of bacteria were determined by spreading ten-fold serial dilutions (1:10 to 1:10^6^) of bacterial cultures on Columbia 5 % sheep blood agar plates (Becton Dickinson), which were incubated at 37 °C for 24 h (*J. psychrophilus* at 24 °C for 72 h) (Table S1).

### Bacteria identification and selection

Bacterial colonies were picked with a sterile toothpick and manually turned ten times in 10 μl distilled water (Invitrogen UltraPure DNase/RNase-free distilled water; Thermofisher Scientific, USA) in a 1.5 ml tube. The tubes were heated at 95 °C for five minutes and then centrifuged for one minute at 13000 rpm. Three μl of supernatant were used for PCR which was performed with 1 μl (10 μM) of forward and reverse primers, 4 μl of MyTaq Red Reaction Buffer (Bioline, UK), 11 μl water and 0.1 μl MyTaq DNA polymerase (Bioline). The PCR cycling conditions, after an initial cycle of denaturation at 95 °C for five minutes, included 40 cycles of denaturation at 95 °C for 20 sec, annealing at 60 °C for 30 sec, extension at 72 °C for 60 sec, followed by a final extension step at 72 °C for seven minutes.

The bacterial specific forward primer 27F (5’ – AGAGTTTGATCCTGGCTCAG - 3’) and the universal reverse primer 805R (5’ – GACTACCAGGGTATCTAATCC – 3) (Tanner et al., 1999) targeting the 16S ribosomal RNA (rRNA) were used. Amplicons were purified using the Qiaquick PCR purification kit (Qiagen, Germany) according to the manufacturer’s instruction, with the exception of using 13 μl of elution buffer instead of 10 μl. DNA concentrations were measured with a microvolume spectrophotometer (Nanodrop One; Thermofisher Scientific) and diluted with deionized water to final concentrations of 7.5 ng/μl. Samples were subsequently sent for Sanger sequencing to a private company (Microsynth, Switzerland). Sequence identities were determined using Nucleotide Basic Local Alignment Search Tool (https://blast.ncbi.nlm.nih.gov/Blast.cgi) utilizing the curated 16S ribosomal RNA sequences database for Bacteria and Archaea. Matrix-Assisted Laser Desorption/Ionization-Time of Flight mass spectrometry (MALDI-TOF MS) was done on individual colonies with low identity (< 99 %) or multi affiliations by a private company (Mabritec AG, Riehen, Switzerland). MALDI-TOF mass spectra were acquired on a microflex Biotyper (Bruker Daltonics, Germany) as described (Cuénod et al., 2021) and the species identified with MABRITECCENTRAL, a database for the comparison and identification of bacterial MALDI-TOF MS spectra (www.mabriteccentral.com) (Mabritec AG). Processed spectra are compared against the SARAMIS™ (Spectral ARchive And <u>Microbial Identification</u> System database, Anagnostec GmbH, 14476 Potsdam, Germany) and PAPMID™ (Putative Assigned Protein Masses for Identification Database, Mabritec AG) databases.

Bacterial species were selected for the behavioural assays based on their 1) harmlessness (biosafety group 1/1+) ensuring a safe future usage as potential skin probiotic, 2) abundance and 3) VIP-scores as determined in our earlier study (Jost et al., 2023). The VIP-score describes the most influential bacteria for the separation between sheep breeds, body parts and origin by PL-SDA (Jost et al., 2023).

### Biting midges

*Culicoides nubeculosus* (Meigen) were laboratory-reared as described (Kaufmann et al., 2015). Female eight-to-ten days old *C. nubeculosus* which did not receive a blood meal for at least four days were used for the behavioural assays.

Field-caught biting midges were collected at a farm at the outskirts of Zürich and at the Zoo Zürich from May until October 2022 for dual-choice behavioural assays and from May until July 2023 for feeding assays using Onderstepoort UV-light suction traps (OVI) (Venter et al., 1996).

Traps were set up around six PM and collected at eight AM the next morning. Biting midges were morphologically identified down to Culicoides group level (Obsoletus, Pulicaris, other) under a microscope (Augot et al., 2017; Meiswinkel and Gomulski, 2004). Captured midges were directly placed into an incubator (24 °C, relative humidity of 85 % and a 16h:8h light:dark cycle) and maintained with ad libitum access to a glucose solution (5 %) provided on cottonwool. Field-caught midges were only used once, between one to six days after collection.

### Dual-choice behavioural assay

A schematic drawing of the constructed Y-tube olfactometer (technical glassblowing workshop Irchel, UZH, Switzerland) for the dual-choice behavioural assays is given in Figure 1. The Y-tube was placed on a white wooden platform at a 30° angle. The test arms were turned downwards when testing *C. nubeculosus* and upwards with field-caught Culicoides, as pilot experiments had shown that this resulted in the highest responses. The temperature and relative humidity in the room were set to 26 °C and 70 %, respectively. An air pump (KNF LABOPORT, Germany) provided a charcoal-filtered, humidified and heated airflow (150 mL/min). A CO_2_-flow activating the insects host-seeking behaviour (Logan et al., 2010) was maintained at 15 mL/min. Both gas flows were combined through Teflon tubes and connected to the two odour adapters containing the test compounds.

**Figure 1:**
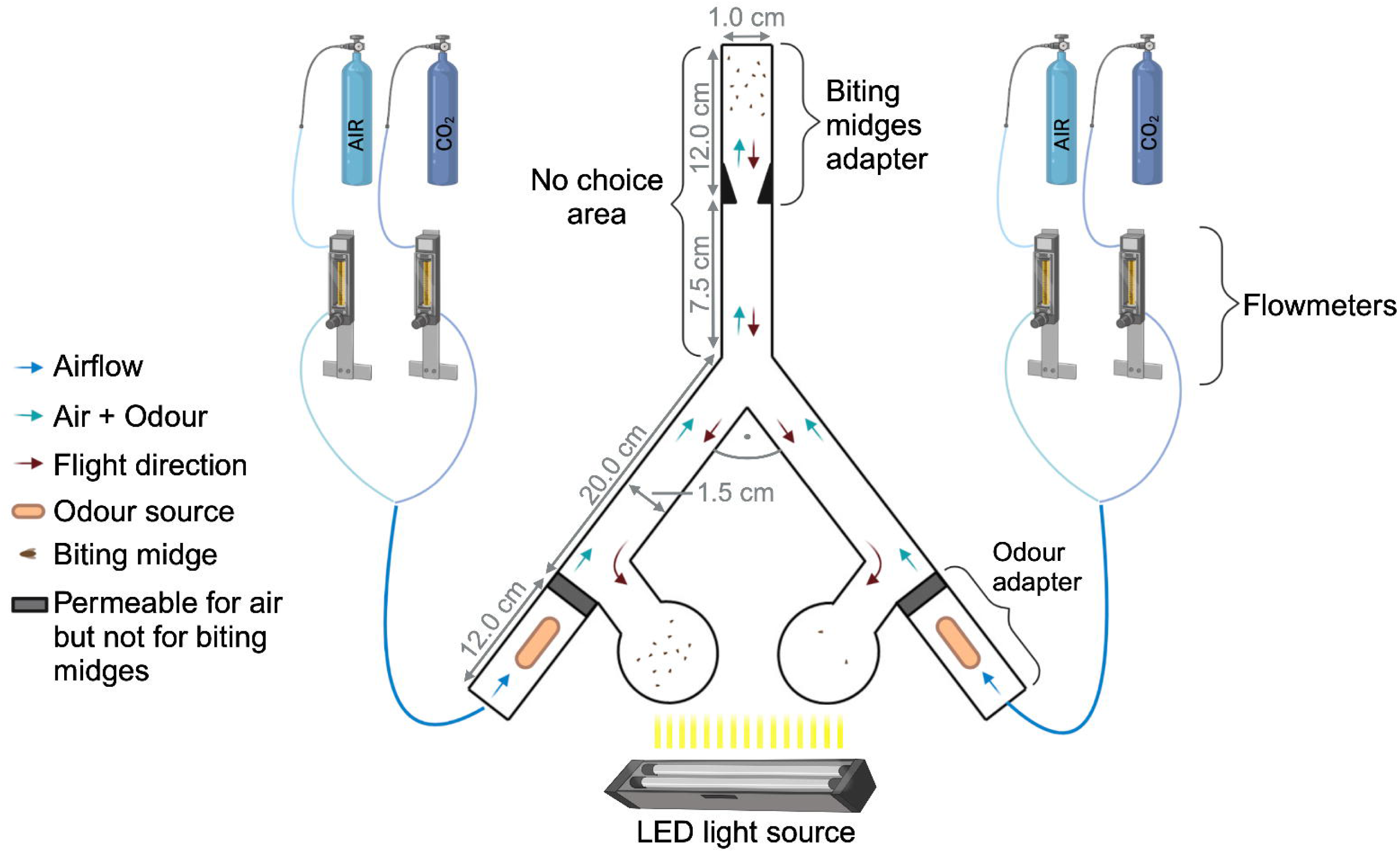
Schematic drawing of the dual-choice Y-tube olfactometer (not to scale). The main tube of the olfactometer (7.5 cm length, Ø 1.5 cm) merges into two test arms of 20.0 cm length and 1.0-1.5 cm diameter, with a 90° angle between them. Each test arm of the Y-tube olfactometer has a glass bulb (50 ml, Duran, Schott AG, Germany) turned to the medial side and a glass odour adapter (12.0 cm length, Ø 1.0 cm) filled with the test compounds on the lateral side. The biting midge’s adapter (12.0 cm length, Ø 0.5-1.0 cm) is inserted at the upper end of the main tube releasing the midges in the olfactometer. Charcoal-filtered, humidified, water bath-heated air (150 mL/min) provided by an air pump and CO_2_ (15 mL/min) and regulated by flowmeters, is directed by Teflon tubes through the odour source to the biting midges. Attracted by LED light, midges move to their preferred odour and are trapped in the glass bulbs, which are removed and frozen for examination of the biting midges. Created with BioRender.com.

One hundred μl of the bacteria cultured in TSB as described above were pipetted onto a filter paper (Ø 25 mm, Macherey-Nagel, Germany) and placed in the glass odour adapters with the help of sterile tweezers. As a control, a filter paper with 100 μl control solution (2050 μl TSB and 50 μl glycerol) was used. Before usage, the filter papers were soaked in acetone and left to dry for 24 h under a fume hood, to minimize intrinsic odours of the filter paper.

The bacterial strains were tested against the control on the experimental days at different timeslot and in randomized order. Six repetitions were performed, with each treatment three times in each arm to avoid side biases. Control experiments with a plain filter paper against a filter paper with 100 μl control solution were also done. In addition, two plain filter papers were tested against each other to detect possible side preferences without odour influences.

Biting midges were transferred to the adapters (Figure 1) via vacuum aspiration. The number of field-caught *Culicoides* spp. used in the trials ranged from 12-37. When testing *C. nubeculosus*, a maximum of 15 individuals were used per adapter to prevent them from clumping together. *Culicoides nubeculosus* from two adapters were released with a time lag of two minutes. Attracted by LED light (Jakob Maul GmbH, Germany, 220-240 V, 50/60 Hz, max. 7 W), midges moved to their preferred odour and are trapped in the glass bulbs, which are removed and frozen for examination of the biting midges. Midges were allowed to make a choice for eight minutes before the position of the individual insects in the olfactometer was recorded. Biting midges that remained in the main tube (‘no choice area’, Figure 1) were considered non-responders and were excluded from further analysis (Isberg et al., 2016).

The olfactometer was washed with pipe cleaners and acetone before and after each experimental day and left to dry under a fume hood for at least 2 h.

### Feeding assay

Female field-caught *Culicoides* spp. (1578 individuals in total, 18-85 individuals per chamber) were fed on membranes as described before by Hochstrasser et al. (2024) with small modifications (Figure 2). In brief: *Culicoides* spp. were starved for 24 hours and transferred into a feeding chamber (Ø 30 mm, height 40 mm) sealed by a feeding membrane (Nescofilm, Bando Chemical IND. LTD.). The inner side of the feeding membrane was coated with 70 μl of bacterial/control treatment and left to dry for 24 h to prevent biting midges getting stuck in the solution. Subsequently, the Culicoides were allowed to blood-feed on six to eight ml anticoagulated cow blood of 27-29 °C for 45 minutes with a room temperature of 24-25 °C and humidity of 55-67 %. The bacterial treatments, the control treatment (medium + glycerol only) and a second control without any treatment were tested simultaneously. After feeding, the Culicoides were frozen at -20 °C and the number of engorged (with red abdomen) and non-engorged individuals counted. Eight replicates of the feeding experiment were performed for each treatment. The feeding chambers were cleaned with water and ethanol after every test. The feeding membranes were discarded after each use.

**Figure 2:**
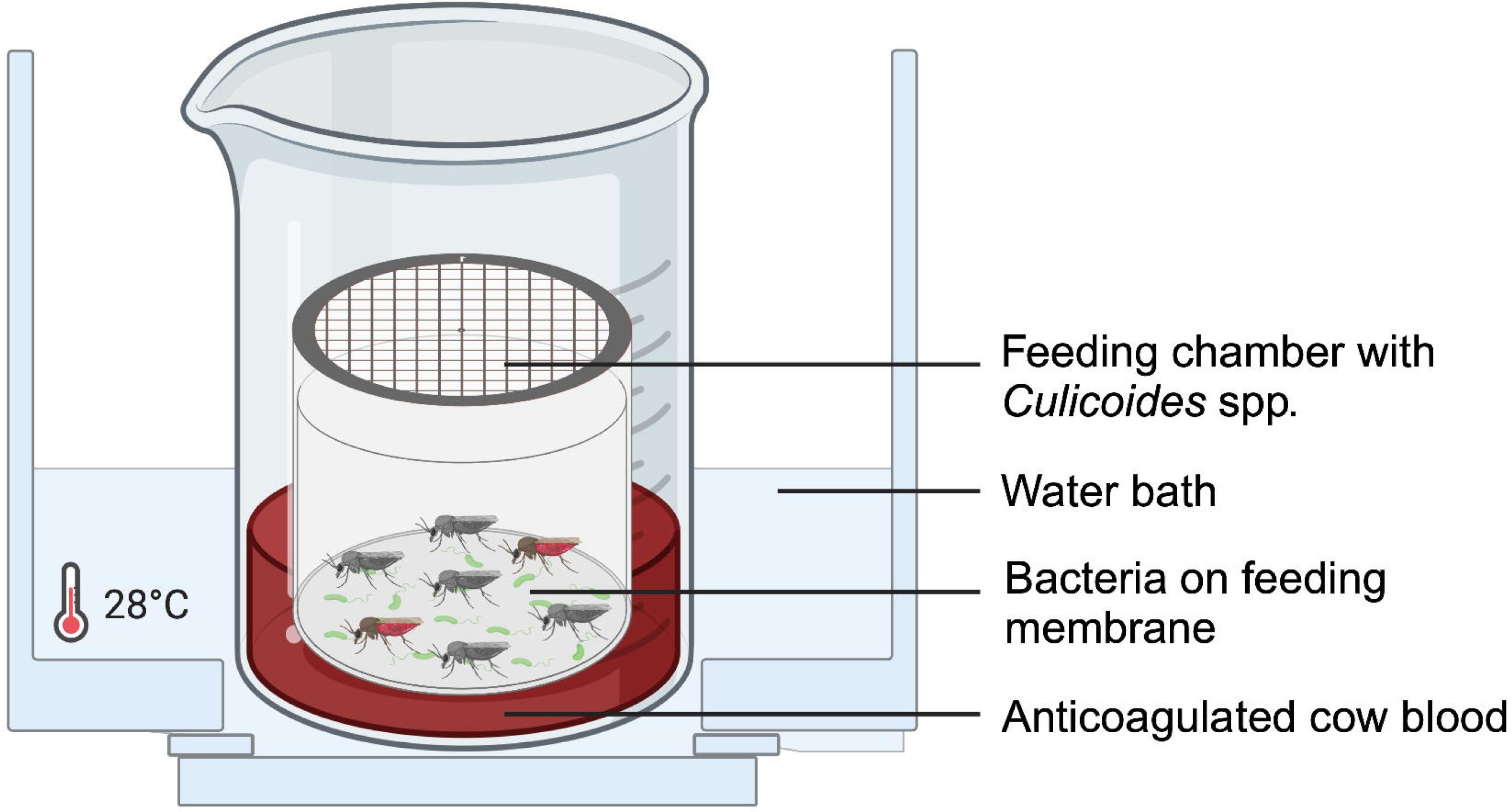
Schematic drawing of the feeding assay. Field-caught Culicoides were inserted in a feeding chamber consisting of a small plastic tube sealed by a feeding membrane coated with bacteria. The feeding chamber was submerged in a beaker filled with cow blood and the Culicoides allowed to blood-feed for 45 minutes. The beaker was placed in a water bath to keep the blood temperature constant between 27-29 °C. Created with BioRender.com.

### Statistical analysis

Chi-squared tests were performed to assess whether the quantities of biting midges in the two olfactometer arms differed (significance threshold P < 0.05) compared to the expected 50:50 distribution.

After checking for no significant homoscedasticity (Levene-Test), a one-way Analysis of Variance (ANOVA) followed by a Tukey’s Honest Significant Difference Test (Tukey’s HSD) was conducted to determine significant differences between treatments used in the membrane feeding assay. All analysis were performed using RStudio and its *stats* package (V. 4.2.2) (R Core Team, 2021) as well as Microsoft Excel (V. 16.77) (Microsoft Corporation, 2018).

## Results

### Skin bacteria

In total, 40 seemingly phenotypically different bacteria colonies were analysed. Thirty-two different species of 20 genera and 11 families were identified by sequencing and/or MALDI-TOF. Twenty bacterial isolates could be identified to species level, one was identified to sister taxa level and eleven to genus level (Table S2). For one isolate, the species identification between MALDI-TOF (*Halalkalibacillus sediminis*) and sequencing (*Micrococcus* sp.) were different and therefore it was decided to exclude this from further analysis because we could not rule out that a contamination had occurred. Based on the selection criteria described, six bacterial species (*Bacillus safensis*, *Bacillus subtilis*, *Micrococcus* sp., *Jeotgalicoccus psychrophilus*, *Staphylococcus equorum* and *Aerococcus urinaeeequi*) were chosen for the behavioural assays. The two most promising bacterial strains (*Micrococcus* sp*., B. safensis*) in the Y-tube olfactometer experiments were further evaluated in a blood feeding assay.

The bacteria used for the behavioural experiments were cultivated in TSB until they reached the stationary phase, as the bacteria in this phase evoked the highest mosquito response (Verhulst et al., 2010). All bacteria reached the stationary phase within 24 h, except *J. psychrophilus* that reached the stationary phase withing 72 h. Growth curves and bacterial concentrations after 24 h (*J. psychrophilus* after 72 h) are given in the supplementary materials (Figure S1, Table S1).

### Biting midges

All field-caught *Culicoides* spp. used for the experiments were identified to group level: 94.2 % (3044/3231) belonged to the Obsoletus group, 5.5 % (177/3231) to the Pulicaris group, and the last 0.3 % (10/3231) to neither of the two (‘others’).

### Dual-choice behavioural assay

The behavioural response of *Culicoides nubeculosus* and field-caught Culicoides towards six bacterial strains was analysed using a Y-tube olfactometer (Figure 3). In cases where over 50 % of the midges that made a choice were located in the treatment arm, the bacteria strain was considered as potentially attractive. Conversely, when more than 50 % were located in the control arm, it indicated a reduced attraction, and the bacteria strain was considered as potentially repellent.

**Figure 3:**
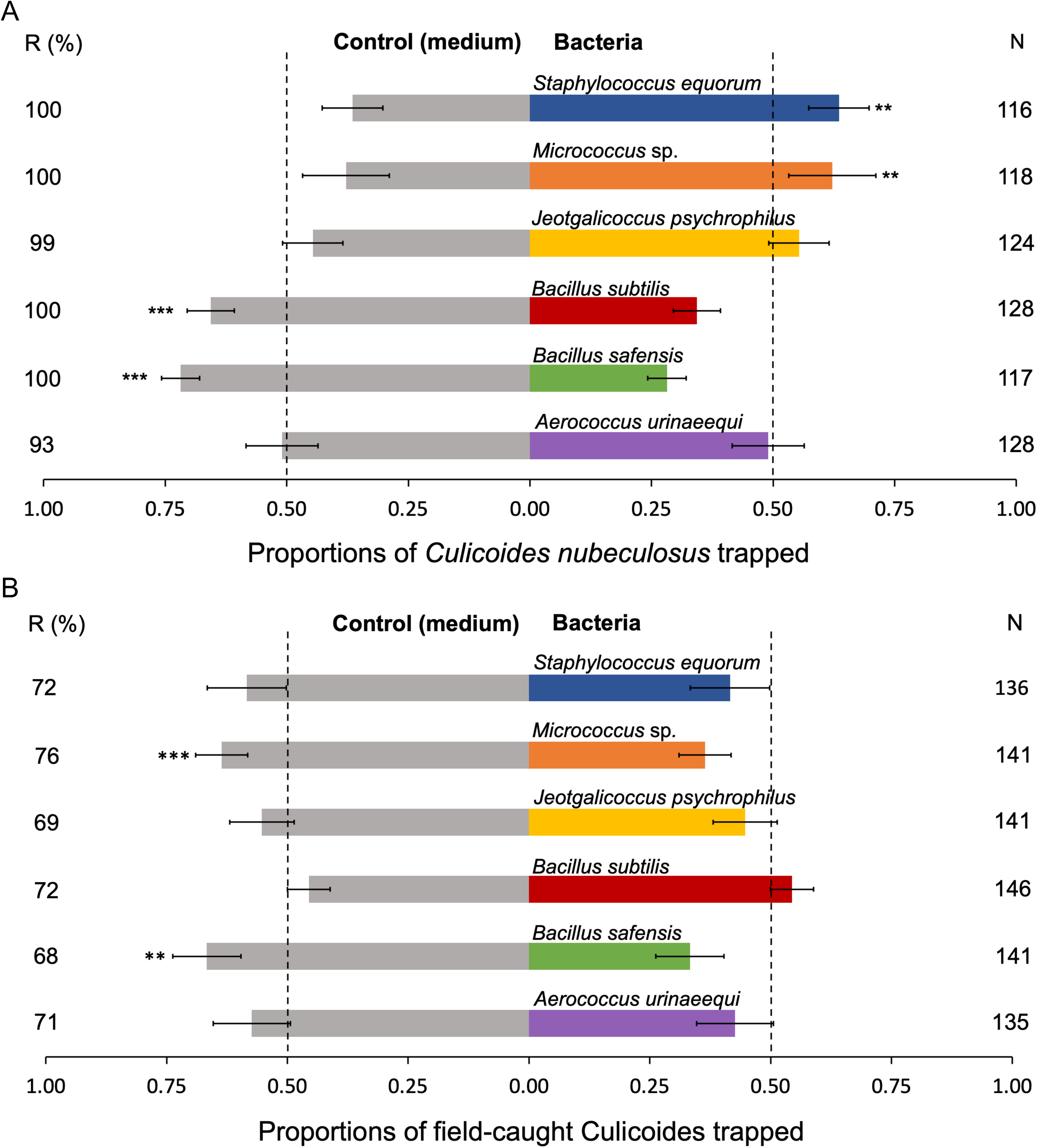
Average response of *Culicoides nubeculosus* (A) and field-caught Culicoides (B) in the dual-choice behavioural assay. Six different bacterial strains (colored bars) were each tested six times against a control (medium (TSB), grey bars). Biting midges were released in the main tube (Figure 1) and allowed to move to their preferred odour source. After eight minutes, the midge’s location was documented, and a proportion of the response calculated. In cases where over 50 % (dashed lines) of the midges that made a choice were located in the treatment arm, the bacteria strain was considered as potentially attractive. Conversely, when more than 50 % were located in the control arm, it indicated a reduced attraction, and the bacteria strain was considered as potentially repellent. Midges that remained in the main tube were considered non-responders and were excluded from the analysis. N: Number of female midges per tested bacteria; R (%): Total average response per tested bacteria. Error bars represent standard errors of the mean; _***_: X^2^-test P < 0.001; _**_: X^2^-test 0.001 ≤ P < 0.01.

From a total of 731 tested female *Culicoides nubeculosus*, 722 (98.8 %) made a choice and were included in further analysis, while of the female field-caught Culicoides (N_total_ = 849) 71.3 % made a choice. *Culicoides nubeculosus* significantly preferred the control side over the bacterial side when testing *B. safensis* (X^2^ = 10.261, d.f. = 1, P ≤ 0.001) and *B. subtili*s (X^2^ = 6.250, d.f. = 1, P ≤ 0.001) (Figure 3A). When testing *Micrococcus* sp. (X^2^ = 3.322, d.f. = 1, P = 0.010) and *S. equorum* (X^2^ = 4.983, d.f. = 1, P = 0.002)*, C. nubeculosus* significantly preferred the bacterial side. Similarly, field-caught Culicoides also showed a significant preference for the control arm when testing *B. safensis* (X^2^ = 3.758, d.f. = 1, P = 0.006) (Figure 3B). In addition, *Micrococcus* sp. (X^2^ = 5.833, d.f. = 1, P ≤ 0.001) was potentially repellent to field-caught biting midges. No bacteria were found that were significantly attractive to field-caught Culicoides. No significant differences were found in the control experiments, confirming that the setup did not have a significant side bias, and the control solution was neither attractive nor repellent (Figure S2).

### Feeding assay

In the feeding assay (Figure 2), in total 1578 field-caught Culicoides were exposed to blood with 4 differently treated membranes. The mean blood feeding rates (proportion of blood-fed Culicoides from a total) showed significant differences between the different treatments (ANOVA, d.f. = 3, P ≤ 0.001) (Figure 4). The highest feeding inhibition was observed when testing the membrane treated with *B. safensis* (5.4 % feeding) compared to the untreated membrane (44.4 %, Tukey’s HSD, P ≤ 0.001), resulting in an 87.9 % feeding reduction. When comparing the feeding rates of *B. safensis* to the control solution membrane (treated with medium + glycerol only) (33.0 %, P ≤ 0.001), a reduction in feeding of 83.8 % was achieved. Feeding rates with *Micrococcus* sp. (23.4 %) were also significantly different to the untreated membrane (P = 0.002) resulting in a 50.4 % feeding reduction, however, this was not significantly different from the control solution (P = 0.274). No significant differences were found when testing the untreated compared to the membrane treated with the control solution (P = 0.157).

**Figure 4:**
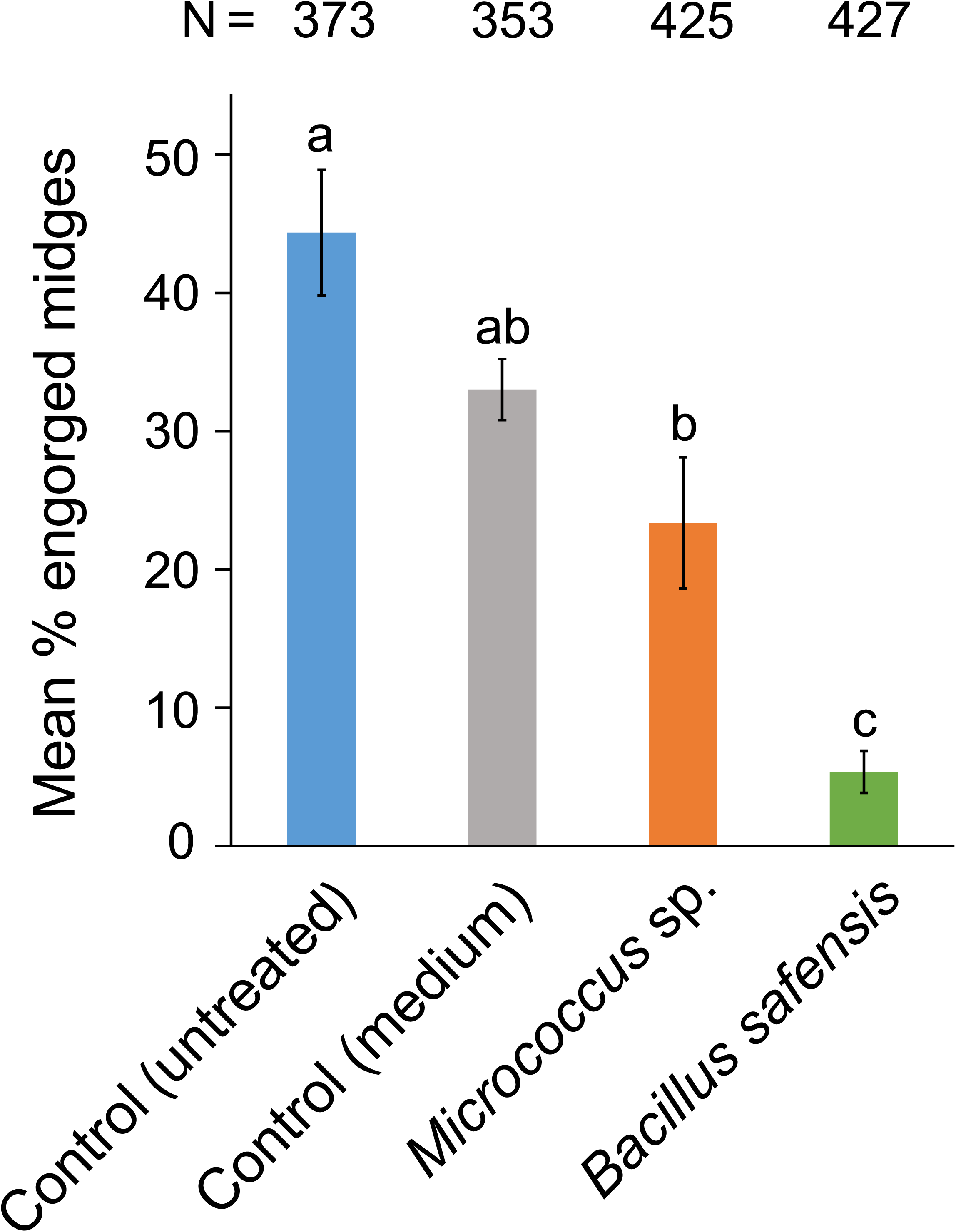
Average feeding rate of field-caught Culicoides in the feeding assay. The trial was performed with four differently treated feeding membranes: Untreated membrane, membrane treated with medium (TSB) only or with bacterial cultures (*Micrococcus* sp., *Bacillus safensis*). Eight replicates of the feeding experiment were performed for each treatment. N: Total number of Culicoides used per treatment. Error bars represent standard errors of the mean. Different superscript letters (a, b, c) indicate statistically significant differences between the treatments (Tukey’s HSD-stest, P ≤ 0.002).

## Discussion

This study aimed to identify sheep skin bacteria that influence the host-selection of biting midges. We show, that isolated sheep skin bacteria evoke both potential attraction and behavioral inhibition in *C. nubeculosus* in a Y-tube olfactometer. Moreover, our research reveals differences in behavioral responses between laboratory-reared and field-caught Culicoides. We observed avoidance behavior towards field-caught Culicoides when exposed to *B. safensis* and *Micrococcus* sp.. In a subsequent blood feeding assay, a feeding reduction of 83.8 % was achieved when testing *B. safensis*.

Skin bacterial isolation from sheep skin showed that the majority of the species belonged to the families Bacillaceae (43.8 %) and Staphylococcaceae (18.8 %). A previous study analyzing the skin bacterial profile of sheep from different breeds, farms and body sites (Jost et al., 2023) by 16S metabarcoding confirmed Staphylococcaceae (relative abundance 18.2 %) and Bacillaceae (relative abundance 2.3 %) as abundant families in the sheep’s skin microbiome. In our bacterial cultures, we identified *Aerococcus* sp., *B. cereus*, *B. pumilus*, *Corynebacterium* sp., *Enterobacter* sp., *Mammaliicoccus scuiri* (former *Staphylococcus scuiri*) *Micrococcu*s sp. and *S. equorum*, mirroring a study that analysed the bacterial skin flora of healthy sheep by phenotypic analysis, MALDI-TOF-MS and 16S rRNA gene sequencing (Haarstad et al., 2014). Interestingly, we also found bacterial species such as *B. thuringiensis* and *Solibacillus* sp. that are primarily associated with the soil showing that sheep skin also carries soil bacteria.

The repellency of six bacterial strains isolated from sheep skin against Culicoides was assessed in a Y-tube olfactometer. We unravelled an avoidance effect towards laboratory-reared *C. nubeculosus* when testing *B. subtilis* and *B. safensis*; the latter also showed comparable effects against field-caught Culicoides (Obsoletus and Pulicaris groups). These field-caught and laboratory-reared biting midges showed opposite behaviour towards *Micrococcus* sp., suggesting a species-specific response. Differences in responses between laboratory-reared and field-caught Culicoides were confirmed in a study investigating their behaviour towards the host-derived volatile nonanal, that was repellent towards *C. nubeculosus* while *C. impunctatus* were attracted (Isberg, 2014). Another study showed lower susceptibility of field-caught Culicoides to deltamethrin exposure than laboratory-reared *C. nubeculosus*, underlining their different responses (De Keyser et al., 2017). Additionally, our study observed different preferences for flight directions in the two species studied: When testing field-caught Culicoides the arms of the Y-tube olfactometer were directed upwards, whereas in *C. nubeculosus* we obtained higher responses with downward-facing arms. These findings imply that the developmental background of animals could play a crucial role in shaping their reactions in laboratory setting, for example due to differences in their own microbiome. However, because the laboratory-reared *C. nubeculosus* were not found in our wild collections it could also be a species effect. This species-specific response emphasizes the importance of working with the insect population of interest as discrepancies in behaviour may arise between field-caught and long-time laboratory-reared individuals (De Keyser et al., 2017).

Interestingly, volatiles produced by *Bacillus* sp. are attractive towards mosquitoes (Verhulst et al., 2010), while we found a potential repellent effect for Culicoides. This highlights the contrast in olfactory behaviour between biting midges and mosquitoes. This behavioural contrast has also been described testing the effects of *d*-allethrin, a commercially available repellent for vector mosquitoes that attracted *C. nubeculosus* (Isberg and Ignell, 2022).

Studies assessing the effects of skin bacteria on mosquitoes have shown that increased microbial diversity reduces the attractiveness of humans (Lucas-Barbosa et al., 2022; Verhulst et al., 2011). Further experiments with different bacterial combinations to increase the microbial diversity could increase the repellent effects. Another study by Zhang et al. (2022) revealed that a flavivirus infection can manipulate host skin microbiome to produce acetophenone, a volatile compound that attracts mosquitoes. Infected mice were bitten more frequently than non-infected ones, which was mediated through the skin microbiome, demonstrating the potential of skin microbial management as control technique against hematophagous insects.

Future research should also focus on the identified attractive bacterial strains (*S. equorum* and *Micrococcus* sp.), as previous studies have shown promising results in reducing the impact of mosquitoes using bacterial compounds as odour baits. (Mweresa et al., 2015; Verhulst et al., 2009). If used together with a repellent, bacterial attractants could be used in a push-pull system (Menger et al., 2014; Wagman et al., 2015), and thereby decrease vector-host interactions more efficiently.

As *B. safensis and Micrococcus* sp. were identified as potentially repellent to field-caught Culicoides in the Y-tube olfactometer, they were chosen for a subsequent feeding assay. Our trials revealed an 83.8 % reduction in blood-fed midges’ when testing a *B. safensis* treated membrane compared to the control solution, which makes *B. safensis* a promising candidate as a novel skin bacterial-based repellent. The employed blood-feeding protocol yielded proportions of engorged insects (44.4 % mean feeding rate) high enough to allow direct comparison of the treatments and their feeding inhibitory effect. Anti-feeding effects of repellents are frequently assessed for mosquitoes (Kajla et al., 2019), yet previous artificial blood-feeding protocols of filed-caught Culicoides were not efficient enough for significant comparisons of feeding deterrents, achieving low engorgement rates between 0.0 % and 7.5 % (Barber et al., 2018). The described protocol (Hochstrasser et al., 2024) has proven to be a valuable tool to evaluate feeding inhibition in field-caught biting midges.

Repellency and attractiveness of host derived-compounds towards Culicoides has been studied and several repellents and attractants were identified (Isberg et al., 2016; Isberg and Ignell, 2022), however, very little is known about the skin bacteria that potentially produce those compounds. Our research narrows this knowledge gap by successfully identifying skin bacterial strains repellent to biting midges. Following steps could be to identify the associated bacterial compounds, for instance with gas chromatography – electroantennographic techniques (Isberg et al., 2016; Qiu et al., 2004), and assessing the efficiency of the repellent bacteria in in vivo settings (Venail et al., 2011; Wichgers Schreur et al., 2021). In addition, it would be interesting to test the efficiency against conventional repellents.

Compared to conventional insecticides that show limited and/or short-lived effectiveness against biting midges (Harrup et al., 2016; Venail et al., 2011), skin probiotics may achieve longer-lasting protection (Lucas-Barbosa et al., 2022), as studies ensure the successful growth of several probiotics under skin-like conditions (Lizardo and Tavaria, 2022). This effect could have even higher potential on animals than in humans as they do not wash and thereby remove the treatment. In addition, animals do not use skincare products and have more body hair, which will favour bacterial growth and sustainability. However, rain or a harsher environment could negatively affect the skin probiotics effectiveness which should be evaluated. Despite several health benefits, some limitations of skin probiotic usage may appear, as the interreference with the skin microbiome may cause allergic reactions and dysbiosis compromising the natural skin barrier. Further studies are necessary to demonstrate the safety of topical probiotic usage.

With the rising prevalence of diseases transmitted by Culicoides (Stokstad, 2023), there is an urgent requirement to devise novel control strategies. Global environmental changes with the inherent potential expansion of the geographical distribution of Culicoides and the prolongation of pathogen transmission periods (Hudson et al., 2023; Sanders et al., 2019) aggravate the current situation of lacking any effective control methods against these insects (Harrup et al., 2016; Shults et al., 2021). Exploitation of skin probiotics against hematophagous insects is still at a very early stage. Building on the results presented, we see potential for skin microbial management as a control technique against biting midges. Next step would be to optimize the application by testing different formulations and bacterial concentrations, which should lead to a first exploratory study in which the probiotic treatment is applied to sheep and the reduction of biting by midges is monitored.

## Acknowledgements

We would like to thank Alexander Mathis for his valuable support and input to the manuscript. Additionally, we would like to thank Alec Hochstrasser for providing the blood feeding protocol and his contribution to the statistical analysis and Laëtitia Cardona (EPFL Lausanne) for the assistance in the genetic bacteria identification. We also thank the Zoo Zürich for the opportunity to perform the insect collections. This work was supported by the Gebert Rüf Stiftung (project no. GRS-089/20). We highly acknowledge the Swiss Federal Food Safety and Veterinary Office as supporter of the Swiss National Centre for Vector Entomology.

## Data availability statement

The data that support the findings of this study are available in Zenodo (https://zenodo.org/doi/10.5281/zenodo.10725750)

The sequences obtained in this study were submitted to the NCBI GenBank and accession numbers (GenBank: PP728454 - PP728484) were assigned.

## CRediT authorship contribution statement

**Paula S. Brok:** Conceptualization, Formal analysis, Investigation, Writing – original

draft. **Stéphanie M. Jost:** Conceptualization, Investigation, Writing – review & editing. **Niels O. Verhulst:** Conceptualization, Funding acquisition, Supervision, Writing – review & editing.

## Declaration of competing interest

The authors declare that they have no known competing financial interests or personal relationships that could have appeared to influence the work reported in this paper.

## Notes

https://zenodo.org/doi/10.5281/zenodo.10725750

### Competing Interest Statement

The authors have declared no competing interest.

