## supplementary materials for "Selection of sheep skin bacteria to reduce blood-feeding by biting midges under laboratory conditions"

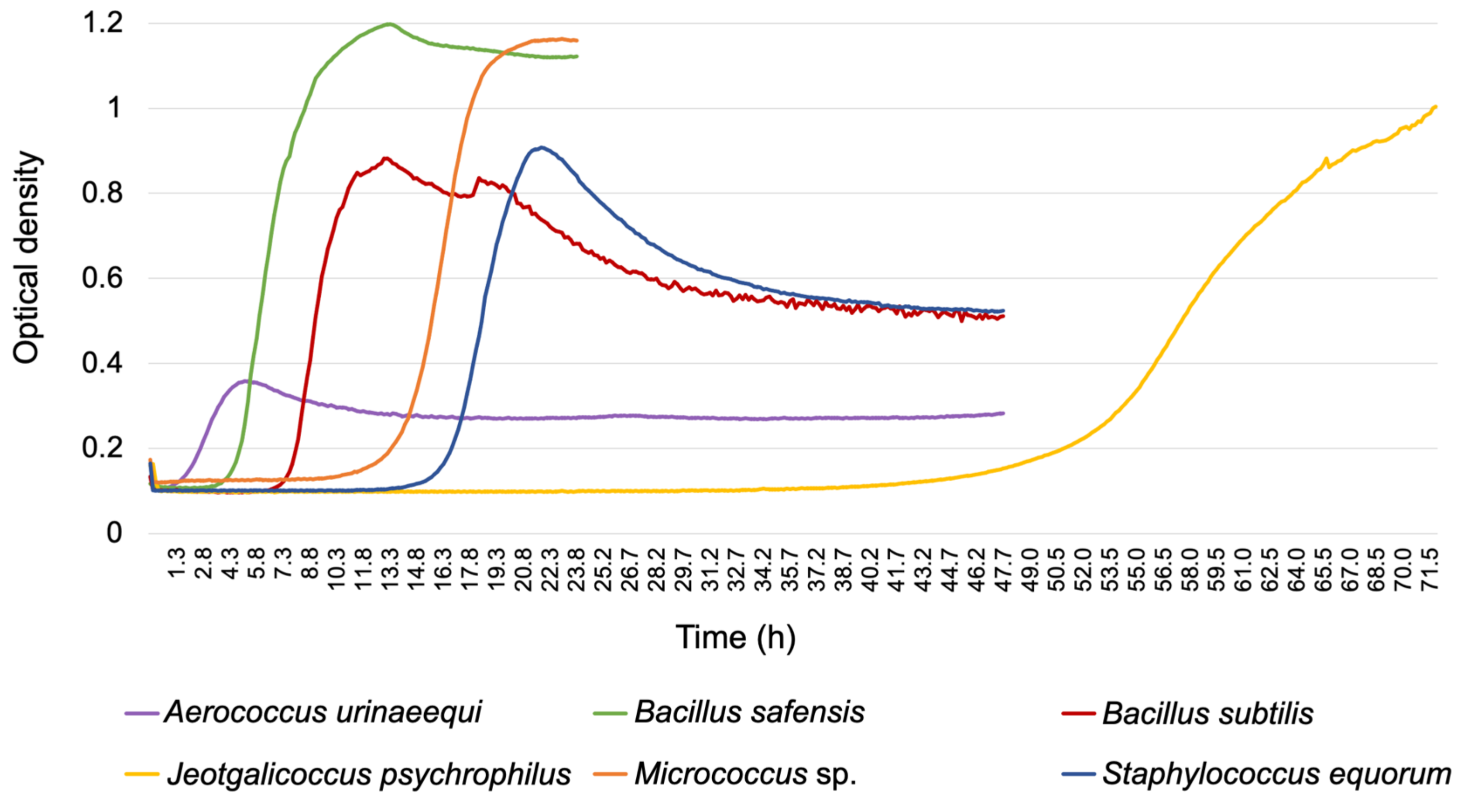


Figure S1: Growth curves measured by the optical density (absorbance 600 nm) over time (h) of the six bacteria species studied (*Aerococcus urinaeequi, Bacillus safensis, Bacillus subtilis, Jeotgalicoccus psychrophilus.*, *Micrococcus* sp., *Staphylococcus equorum*) at 37 °C (*Jeaotgalicoccus psychrophilus* 24 °C).

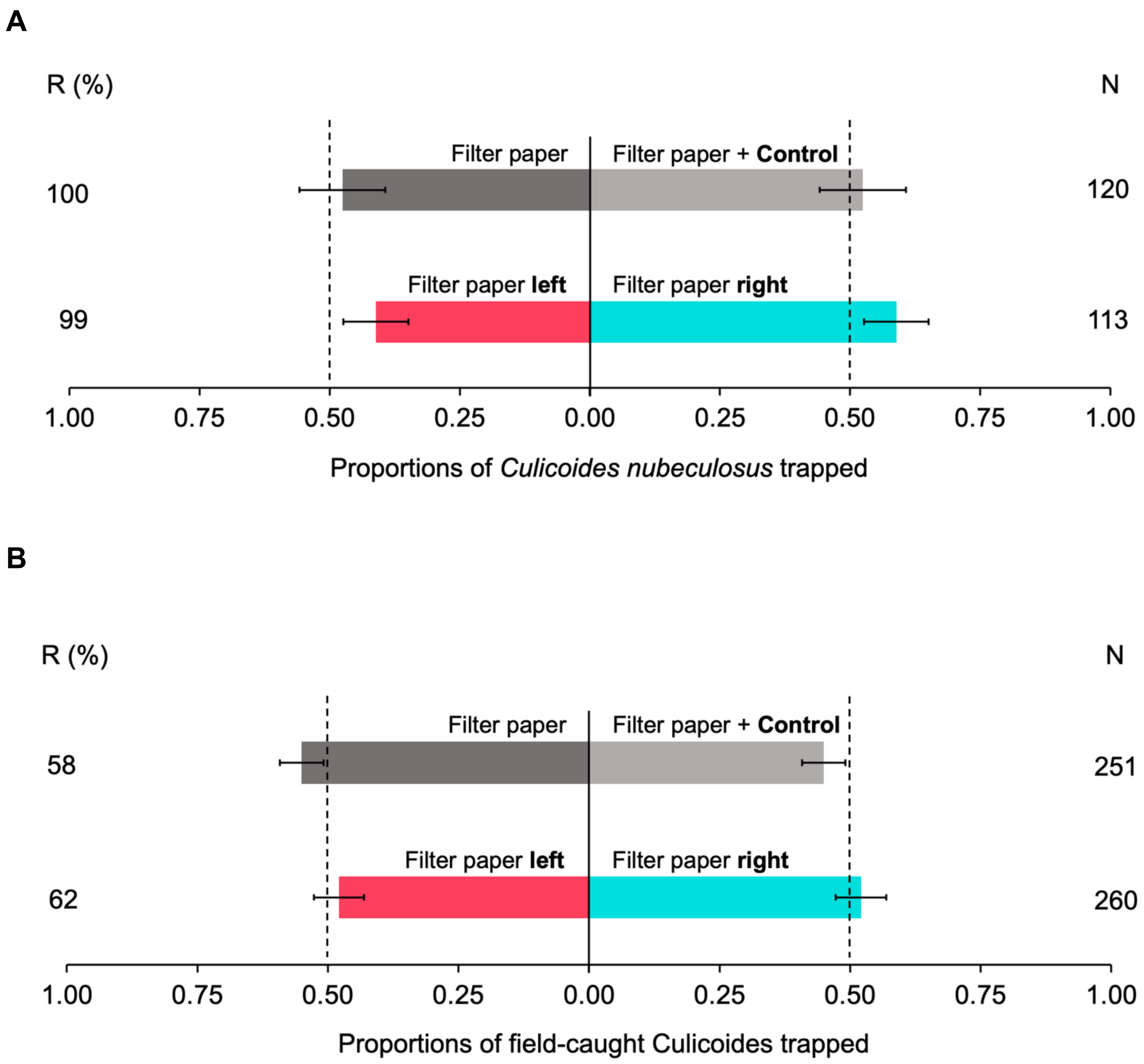


Figure S2: Control experiments for *Culicoides nubeculosus* (A) and field-caught Culicoides (B) in the dual-choice behavioural assay. The control solution (2050 µl TSB and 50 µl glycerol) on filter paper was tested against the filter paper to detect any behavioural effect of the control solution. Plain filter papers were compared to detect possible side preferences without odour influences. N: Number of female midges per tested bacteria; R (%): Total average response. Error bars represent standard errors of the mean. No significant differences were found.

Table S1: Bacterial concentrations used for experiments. Concentrations were determined by counting the colony forming units (CFU) in a dilution series.

| Bacteria | Incubation time (h) | Incubation temperature (°C) | Dilution | CFU | Concentration (CFU/µl) |
|---|---|---|---|---|---|
| *Aerococcus urinaeequi* | 24 | 37 | $10^{-5}$ | 84 | 84000 |
| *Bacillus safensis* | 24 | 37 | $10^{-5}$ | 257 | 257000 |
| *Bacillus subtilis* | 24 | 37 | $10^{-6}$ | 25 | 250000 |
| *Jeotgalicoccus psychrophilus* | 72 | 24 | $10^{-6}$ | 57 | 570000 |
| *Micrococcus* sp. | 24 | 37 | $10^{-6}$ | 266 | 2660000 |
| *Staphylococcus equorum* | 24 | 37 | $10^{-6}$ | 50 | 500000 |

Table S2: Identification of skin bacterial samples. The sequence ID is describing the top BLAST (NCBI) results for bacterial 16S rRNA sequences. MALDI-TOF was performed on individual colonies with identity percentages of less than 99 % or in cases of multi affiliations. The obtained sequences were submitted to the NCBI GenBank and accession numbers (GenBank: PP728454 - PP728484) were assigned.

| Isolate no. | GenBank accession no. | Sequence ID (Top BLAST hits) | Query coverage | Sequence identity | MALDI-TOF ID | **Identified species** | Genus | Family (prevalence) |
|---|---|---|---|---|---|---|---|---|
| 1 | PP728454 | *Aerococcus viridans* | 100.00% | 99.80% | *Aerococcus urinaeequi* | ***Aerococcus urinaeequi**** | Aerococcus | Aerococcaceae (6.25%) |
| | | *Aerococcus urinaeequi* | 100.00% | 99.80% | | | | |
| 2 | PP728455 | *Aerococcus urinaeequi* | 100.00% | 99.79% | n.d. | ***Aerococcus* sp.** | | |
| | | *Aerococcus viridans* | 100.00% | 99.37% | | | | |
| 3 | PP728456 | *Bacillus altitudinis* | 100.00% | 100.00% | *Bacillus altitudinis* | ***Bacillus altitudinis*** | Bacillus | Bacillaceae (43.75%) |
| | | *Bacillus australimaris* | 100.00% | 100.00% | | | | |
| | | *Bacillus xiamenensis* | 100.00% | 100.00% | | | | |
| 4 | PP728457 | *Bacillus licheniformis* | 100.00% | 99.53% | *Bacillus glycinifermentans* | ***Bacillus glycinifermentans*** | | |
| | | *Bacillus sonorensis* | 100.00% | 99.53% | | | | |
| | | *Bacillus aerius* | 100.00% | 99.53% | | | | |
| 5 | PP728458 | *Bacillus pumilus* | 100.00% | 99.80% | *Bacillus pumilus* | ***Bacillus pumilus*** | | |
| | | *Bacillus safensis* | 100.00% | 99.80% | | | | |
| 6 | PP728459 | *Bacillus safensis* | 100.00% | 99.80% | *Bacillus safenis* | ***Bacillus safensis**** | | |
| | | *Bacillus pumilus* | 100.00% | 99.80% | | | | |
| 7 | PP728460 | *Bacillus subtilis* | 100.00% | 99.97% | n.d. | ***Bacillus* sp.** | | |
| | | *Bacillus inaquosorum* | 100.00% | 99.48% | | | | |
| | | *Bacillus mojavensis* | 100.00% | 99.48% | | | | |
| | | *Bacillus tequilensis* | 100.00% | 99.48% | | | | |
| 8 | PP728461 | *Bacillus tequilensis* | 100.00% | 100.00% | *Bacillus subtilis* | ***Bacillus subtilis**** | | |
| | | *Bacillus subtilis* | 100.00% | 99.40% | | | | |
| 9 | PP728462 | *Bacillus thuringiensis* | 100.00% | 100.00% | *Bacillus thuringiensis* | ***Bacillus thuringiensis*** | | |
| | | *Bacillus proteolyticus* | 100.00% | 100.00% | | | | |
| | | *Bacillus wiedmannii* | 100.00% | 100.00% | | | | |
| | | *Bacillus toyonensis* | 100.00% | 100.00% | | | | |
| 10 | PP728463 | *Bacillus mobilis* | 100.00% | 100.00% | *Bacillus wiedmannii* | ***Bacillus wiedmannii*** | | |
| | | *Bacillus wiedmannii* | 100.00% | 99.75% | | | | |
| | | *Bacillus proteolyticus* | 100.00% | 99.75% | | | | |
| | | *Bacillus thuringiensis* | 100.00% | 99.75% | | | | |
| 11 | PP728464 | *Lysinibacillus parviboronicapiens* | 100.00% | 98.43% | *Lysinibacillus sp.* | ***Lysinibacillus* sp.** | Lysinibacillus | |
| | | *Lysinibacillus cresolivorans* | 94.00% | 96.67% | | | | |
| | | *Lysinibacillus macroides* | 94.00% | 95.83% | | | | |
| 12 | PP728465 | *Peribacillus simplex* | 100.00% | 100.00% | *Perbacillus frigoritolerans* | ***Peribacillus frigoritolerans*** | Peribacillus | |
| | | *Peribacillus frigoritolerans* | 100.00% | 99.78% | | | | |
| 13 | PP728466 | *Peribacillus muralis* | 100.00% | 97.56% | *Peribacillus muralis* | ***Peribacillus muralis*** | | |
| | | *Peribacillus simplex* | 100.00% | 97.56% | | | | |
| | | *Peribacillus frigoritolerans* | 100.00% | 97.56% | | | | |
| 14 | PP728467 | *Peribacillus simplex* | 100.00% | 99.56% | *Peribacillus simplex* | ***Peribacillus simplex*** | | |
| | | *Peribacillus frigoritolerans* | 100.00% | 99.56% | | | | |
| 15 | PP728468 | *Priestia aryabhattai* | 100.00% | 100.00% | *Priestia aryabhattai/megaterium* | ***Priestia aryabhattai/megaterium*** | Priestia | |
| | | *Priestia megaterium* | 100.00% | 100.00% | | | | |
| | | *Bacillus zanthoxyli* | 100.00% | 100.00% | | | | |
| 16 | PP728469 | *Psychrobacillus lasiicapitis* | 100.00% | 99.12% | n.d. | ***Psychrobacillus* sp.** | Psychrobacillus | |
| | | *Psychrobacillus insolitus* | 100.00% | 98.46% | | | | |
| | | *Psychrobacillus psychrodurans* | 100.00% | 98.25% | | | | |
| 17 | PP728470 | *Corynebacterium casei* | 100.00% | 100.00% | n.d. | ***Corynebacterium casei*** | Corynebacterium | Corynebacteriaceae (3.13%) |
| 18 | PP728471 | *Enterococcus casseliflavus* | 100.00% | 100.00% | *Enterococcus casseliflavus* | ***Enterococcus casseliflavus*** | Enterococcus | Enterococcaceae (3.13%) |
| 19 | PP728472 | *Curtobacterium flaccumfaciens* | 100.00% | 100.00% | n.d. | ***Curtobacterium flaccumfaciens*** | Curtobacterium | Microbacteriaceae (3.13%) |
| 20 | PP728473 | *Arthrobacter koreensis* | 100.00% | 98.61% | n.d. | ***Arthrobacter* sp.** | Arthrobacter | Micrococcaceae (6.25%) |
| | | *Arthrobacter gandavensis* | 100.00% | 98.61% | | | | |
| | | *Arthrobacter luteolus* | 100.00% | 98.61% | | | | |
| 21 | PP728474 | *Kocuria atrinae* | 100.00% | 99.52% | n.d. | ***Kocuria* sp.** | Kocuria | |
| | | *Kocuria carniphilia* | 100.00% | 99.35% | | | | |
| 22 | PP728475 | *Acinetobacter lwoffii* | 100.00% | 99.57% | n.d. | ***Acinetobacter lwoffii*** | Acinetobacter | Moraxellaceae (6.25%) |
| 23 | PP728476 | *Acinetobacter baumannii* | 100.00% | 99.72% | n.d. | ***Acinetobacter baumannii*** | | |
| 24 | PP728477 | *Rhodococcus coprophilus* | 100.00% | 99.05% | *Rhodococcus coprophilus* | ***Rhodococcus coprophilus*** | Rhodococcus | Nocardiaceae (3.13%) |
| 25 | PP728478 | *Paenibacillus mobilis* | 100.00% | 100.00% | n.d. | ***Paenibacillus* sp.** | Paenibacillus | Paenibacillaceae (3.13%) |
| | | *Paenibacillus xylanexedens* | 100.00% | 100.00% | | | | |
| | | *Paenibacillus oceanisediminis* | 100.00% | 100.00% | | | | |
| 26 | PP728479 | *Solibacillus silvestris* | 100.00% | 99.54% | n.d. | ***Solibacillus* sp.** | Solibacillus | Planococcaceae (3.13%) |
| | | *Solibacillus kalamii* | 100.00% | 99.31% | | | | |
| | | *Solibacillus isronensis* | 100.00% | 99.08% | | | | |
| 27 | PP728480 | *Jeotgalicoccus psychrophilus* | 100.00% | 100.00% | *Jeotgalicoccus psychrophilus* | ***Jeotgalicoccus psychrophilus**** | Jeotgalicoccus | Staphylococcaceae (18.75%) |
| | | *Jeotgalicoccus aerolatus* | 100.00% | 100.00% | | | | |
| | | *Jeotgalicoccus nanhaiensis* | 100.00% | 100.00% | | | | |
| | | *Jeotgalicoccus halotolerans* | 100.00% | 100.00% | | | | |
| 28 | PP728481 | *Macrococcus canis* | 100.00% | 98.73% | *Macrococcus sp.* | ***Macrococcus* sp.** | Macrococcus | |
| | | *Macrocuccus caseolyticus* | 100.00% | 98.52% | | | | |
| 29 | PP728482 | *Mammaliicoccus vitulinus* | 100.00% | 99.78% | n.d. | ***Mammaliicoccus* sp.** | Mammaliicoccus | |
| | | *Mammaliicoccus fleuretti* | 100.00% | 99.11% | | | | |
| 30 | (<150 bp)** | *Mammaliicoccus sciuri* | 100.00% | 92.19% | *Mammaliicoccus sciuri* | ***Mammaliicoccus sciuri*** | | |
| 31 | PP728483 | *Micrococcus luteus* | 100.00% | 100.00% | *unidentifiable spectrum* | ***Micrococcus* sp.*** | Micrococcus | |
| | | *Micrococcus aloeverae* | 100.00% | 100.00% | | | | |
| | | *Micrococcus yunnanensis* | 100.00% | 100.00% | | | | |
| | | *Micrococcus cohnii* | 100.00% | 100.00% | | | | |
| | | *Micrococcus endophyticus* | 100.00% | 100.00% | | | | |
| | | *Arthrobacter echini* | 100.00% | 100.00% | | | | |
| | | *Micrococcus antarcticus* | 100.00% | 100.00% | | | | |
| 32 | PP728484 | *Staphylococcus equorum* | 100.00% | 99.79% | *Staphylococcus equorum* | ***Staphylococcus equorum*** | Staphylococcus | |

* used in experiments

** Short sequences (<150 bp) are automatically excluded from GenBank submission.
